# Enhancing biochemical resolution by hyper-dimensional imaging microscopy

**DOI:** 10.1101/431452

**Authors:** Alessandro Esposito, Ashok R. Venkitaraman

## Abstract

Two decades of high-paced innovation have improved the spatial resolution of fluorescence microscopy to enable molecular resolution combined with the low-invasiveness and specificity characteristic of optical microscopy. However, fluorescence microscopy also enables scientists and clinicians to map and quantitate the physico-chemical properties (*e.g.*, analyte concentration, enzymatic activities and protein-protein interactions) of biological samples. But the optimization of the biochemical resolving power in fluorescence microscopy is not as well-developed compared to its spatial resolution. Typical techniques rely on the observation of individual properties of fluorescence thus limiting the opportunities for sensing and multiplexing. Aiming to overcome existing limitations, we demonstrate a new imaging paradigm — Hyper Dimensional Imaging Microscopy (HDIM) — that enables the orthogonal properties of fluorescence emission (excited state lifetime, polarization and spectra) in biological samples to be quantified simultaneously and efficiently. Therefore, akin to how multi-dimensional separation in mass-spectroscopy and multi-dimensional spectra in NMR impacted proteomics and structural biology, we envisage that HDIM spectra of unprecedented dimensionality will impact the fields of systems biology and medical diagnostics by maximizing the biochemical resolving power of fluorescence microscopy.

## Introduction

The biochemical environment of tissues and cells can be probed either by analysing the fluorescence of several naturally occurring, often metabolic-related, biomolecules (*e.g.*, various forms of nicotinamide adenine dinucleotide (NAD^+^) and flavin adenine dinucleotide)^1,2^ or by analysing the fluorescence of environmentally sensitive fluorophores (*e.g.*, organic molecules and fluorescent proteins sensitive to pH) introduced into the sample by chemical or genetic means^3,4^. FRET (Foerster resonance energy transfer) is also a well-established and widely-used technique that enables cellular metabolism (*e.g.*, with glucose and ATP probes^5,6^) and signalling (*e.g.*, with phosphorylation, acetylation and methylation probes^7^) to be mapped on single living cells. Often, these assays alter several properties of fluorescence. For instance, the heterogeneous biochemical milieu of tissues introduces complex optical-biochemical signatures into a specimen’s fluorescence, or FRET alters spectra, lifetime and polarization of the FRET pair emission. However, state-of-the-art biochemical imaging techniques often rely just on the detection of simple optical features. We hypothesized that the simultaneous detection of multiple characteristics of fluorescence would permit to extend significantly the biochemical resolving power in fluorescence microscopy, thus supporting more precise measurements or increased multiplexing capabilities (*e.g.*, multiple diagnostic markers or biochemical probes). Here, we illustrate the implementation of a novel detection paradigm that enables the parallel detection of all properties of fluorescence (“*hyper dimensional detection*”, see **Supp. Note 1**) and provide the first analytical tools (*HDIM-toolbox;* **See Supp. Note 2**) to handle such complex datasets. We demonstrate how *Hyper Dimensional Imaging Microscopy* (HDIM) maximizes the biochemical resolving power of fluorescence microscopy and provides simple proof-of-concept experiments to illustrate how the increased resolution and multiplexed capabilities could be applied for biomedical and clinical applications.

## Results

### Improving biochemical resolving power

HDIM requires a detection system with multiple parallel detection channels with electronics that generate histograms containing information about the polarization, colour and arrival time of each detected photon at every position within the sample. The process of partitioning photons into “detection channels” of specific spectroscopic meaning tends to maximize information on the optical-biochemical system under investigation. Moreover, not using analysers or filters, HDIM minimizes photon losses therefore avoiding the increase of the already long exposure times required by time-resolved techniques. The formal description of the Fisher information for hyper dimensional detection permits to demonstrate the net increase in the biochemical resolving power that HDIM could provide^8^. In **Supp. Fig. 1 and Supp. Note 2**, we provide a brief description of the theory and a simple graphical interpretation of the theoretical results.

**Fig. 1a** and **Supp. Figs. 2-3** depict the experimental setup and the conceptual representation of HDIM. A pulsed laser (here a Ti:Sapphire laser for two-photon excitation) provides tightly controlled excitation light of known timing, polarization and spectra that is delivered to the sample with a laser scanning microscope. Fluorophores and the biochemical environment of the sample reshape the excitation signal introducing complex optical signatures in the emitted fluorescence that is fully characterized by hyper-dimensional detection achieved with a pair of multi-wavelength time-correlated single photon-counting detectors (**Methods**). The photophysics (biochemistry) of the sample is thus described by 2,048 independent values, *i.e.* photon-counts accumulated into 16 spectral bins, 2 polarization states and 64 time-gates within each pixel of the acquired image. With the use of HDIM-tailored analysis algorithms, it is then possible to retrieve biochemical signatures from the complete photophysical characterization of the sample.

**Figure 1.**
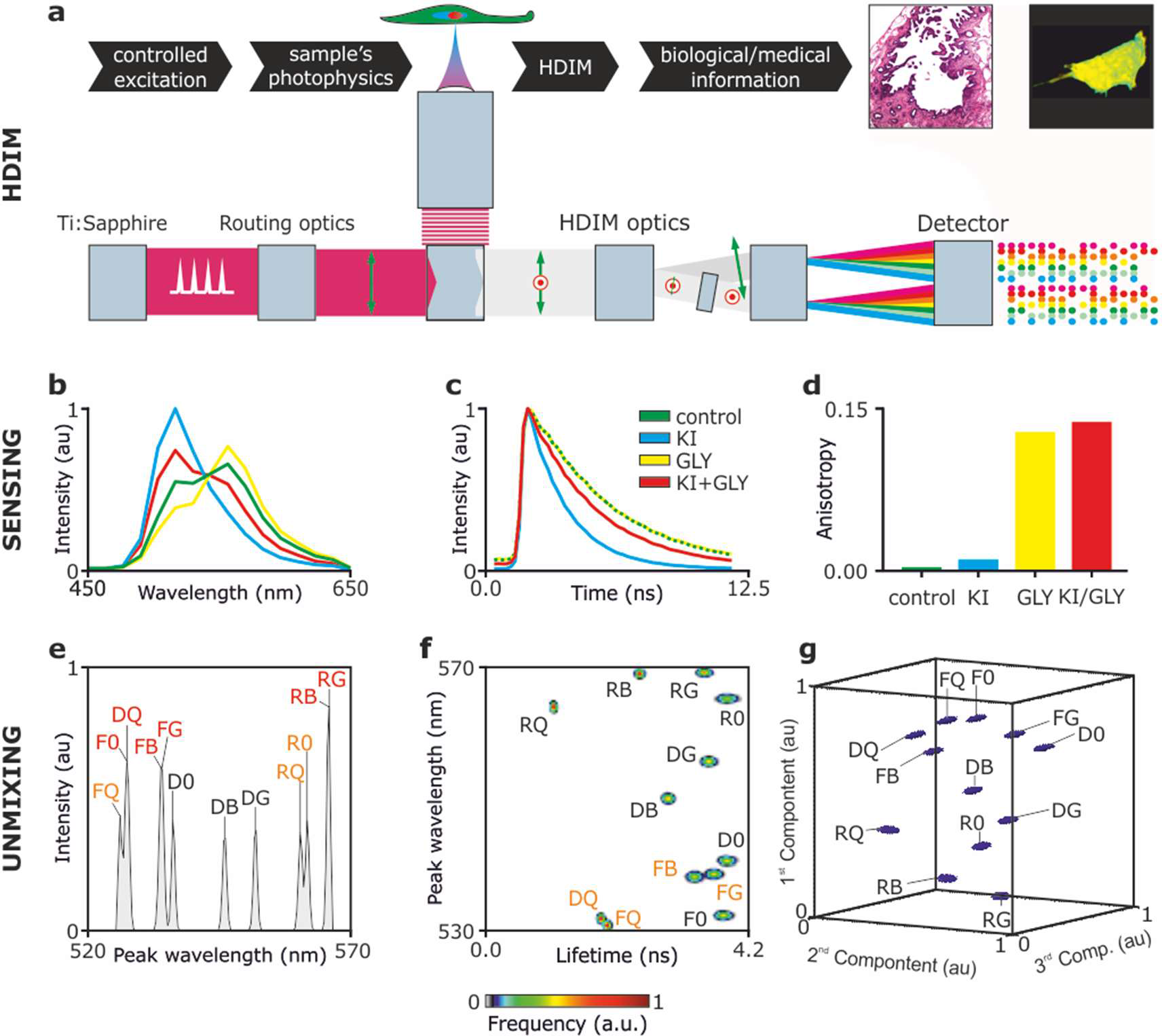
Sensing and unmixing by hyper-dimensional imaging microscopy (HDIM) HDIM relies on the controlled excitation of the sample and by the analysis of biochemical signatures introduced into the photophysical properties of fluorophores within the sample (a). Sensing the biochemical environment of fluorophores (here R6G and FITC) can be achieved with the simultaneous detection of emission spectra (b), fluorescence lifetime (c) and anisotropy (d). Unmixing of different biochemical environment sensed by R6G and FITC is enhanced by the increased dimensionality of the analysed photophysical signatures: spectral peak (e), spectral peak vs lifetime (f) and three-dimensional PCA (g). Peaks are marked with the labels “F” for FITC, “R” for R6G and “D” for a R6G and FITC mixture, followed by “0” (0% glycerol, 200 μM KCl), G (65% glycerol, 200 μM KCl), Q (0% glycerol, 100 μM KCl, 100 μM KCl) or B (65% glycerol, 100 μM KCl, 100μM KI). Red and orange labels indicate non-resolved and partially overlapping peaks, respectively. Excitation: 840nm.

To demonstrate the sensing and un-mixing capabilities of HDIM, we imaged various solutions prepared with 1 μM rhodamine 6G and 10 μM fluorescein. Glycerol was used to reduce the rotational freedom of the fluorophores and equimolar substitution of potassium chloride with the quencher potassium iodide was employed to reduce their fluorescence lifetime (see **Supp. Figs. 3-5** for the complete dataset). **Fig. 1b-d** shows how the emission spectra, lifetime and anisotropy of a mixture of rhodamine 6G and fluorescein are altered by their biochemical environment and how HDIM can sense these changes. By means of spectra of increasing dimensionality, HDIM can successfully separate different physico-chemical environments. **Supp. Fig. 4a-f** further demonstrates that the twelve different mixtures of rhodamine 6G, fluorescein, glycerol and potassium iodide can be resolved only with spectra of higher dimensionality compared to typical one-dimensional spectral information. We tested also the benefits of implementing Principal Component Analysis (PCA) as mean of contrast-enhancement for the analysis of HDIM datasets (see **Fig. 1g** and **Supp. Fig. 5g-l**). PCA increased the minimal observable difference among photophysical signatures within the sample, *i.e.* further improved the (biochemical) resolution of fluorescence microscopy.

### Contrast enhancement during post-processing

We then investigated whether the increased resolving power of HDIM could reveal structures otherwise not or poorly visible when sensing individual optical properties. To test this possibility, we acquired images of *Convallaria majalis* stained with Safranin and Fast Green, a typical sample used to test microscopy techniques. **Fig. 2** illustrates the wealth of information that it is acquired by HDIM. In this case, the dataset is excitation-resolved as well by scanning the Ti:Sapphire laser from 750nm to 1000nm in steps of 50nm. We demonstrate how fluorescence lifetime, anisotropy and emission spectra change as function of excitation wavelength. The complexity of the optical signatures acquired by HDIM is shown in **Fig. 2b** as hyper-dimensional spectral signature or HDSS.

**Figure 2.**
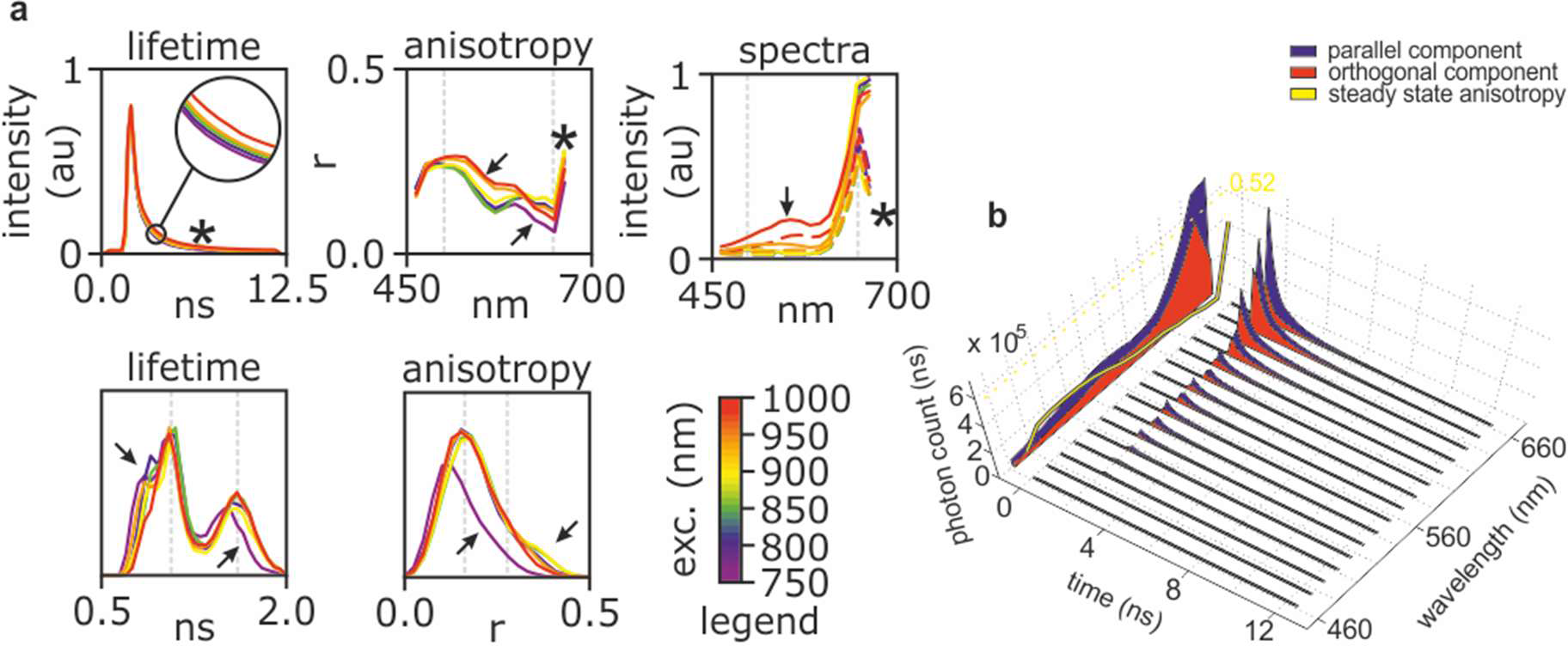
Sensing by hyper-dimensional imaging microscopy. (**a**) Fluorescence lifetime decays (top-left), anisotropy spectra (top-center) and emission spectra (top-right) as a function of excitation wavelength measured on a single field of view of *Convallaria majalis*. The bottom panels show excitation-dependent distributions of fluorescence lifetimes (bottom-left) and anisotropies (bottom-right). Arrows and star highlight correlated features that are modulated by excitation wavelength. (**b**) Example of hyper dimensional spectral signature (HDSS) at 800nm excitation wavelength. Time-decays are shown for its spectral and polarization components. On the back projection of the 3D plot, polarization-dependent spectra and anisotropy spectrum are shown, with the dashed yellow line marking the maximum of fluorescence anisotropy of 0.52 that can be measured with TPE.

**Fig. 3a** shows an intensity image of the sample excited at 800nm. The specific optical properties of the sample can be mapped to two-dimensional maps through simple projections of the abstract 2,048-multidimensional space where each pixel can be described. Projections can be either based on physical quantities (*e.g.*, fluorescence anisotropy), statistical quantities (*e.g.*, PCA or non-negative matrix factorization) or perception-based features (*e.g.*, true colour). The latter is exemplified in **Fig. 3b** which is showing a red-green-blue (RGB) composite image of the specimen as if it was observed through the eyepieces by naked eye. To achieve this representation, first the 2,048-dimensional HDIM dataset is projected on a spectral-only space, effectively summing all photons in each individual time-and polarization-bins. Subsequently, photons from each spectral bin is weighted accordingly to an eye-sensitivity matrix and summed up into three colour channels. Similarly, representation of physical quantities can be synthesized by projecting the HDIM data on other dimensions without applying any weighing factor; for instance, **Fig. 3c,d** show synthetic fluorescence lifetime (FLIM) and anisotropy (FAIM) images (see also **Supp. Fig 7**) generated by projecting HDIM datasets onto the relevant dimensions. To assess if the increased photophysical/biochemical resolution of HDIM translates into contrast enhancement (**Fig. 1g**), we also project the HDIM dataset to an RGB composite showing the first three principal components (**Fig. 3e-h** and **Supp. Fig. X**). PCA is agnostic about the composition of the sample and it merely enhances the contrast for each and between each component. In fact, **Fig. 3e** shows structures of *Convallaria majalis* that colour, FLIM and FAIM images do not highlight. From the inspection of the individual principal components (**Fig. 3f-h**), it is possible to establish that Safranin and Fast Green are detected as first and second principal components, respectively. Autofluorescence is loaded into the third principal component thus providing an additional mean of contrast.

**Figure 3.**
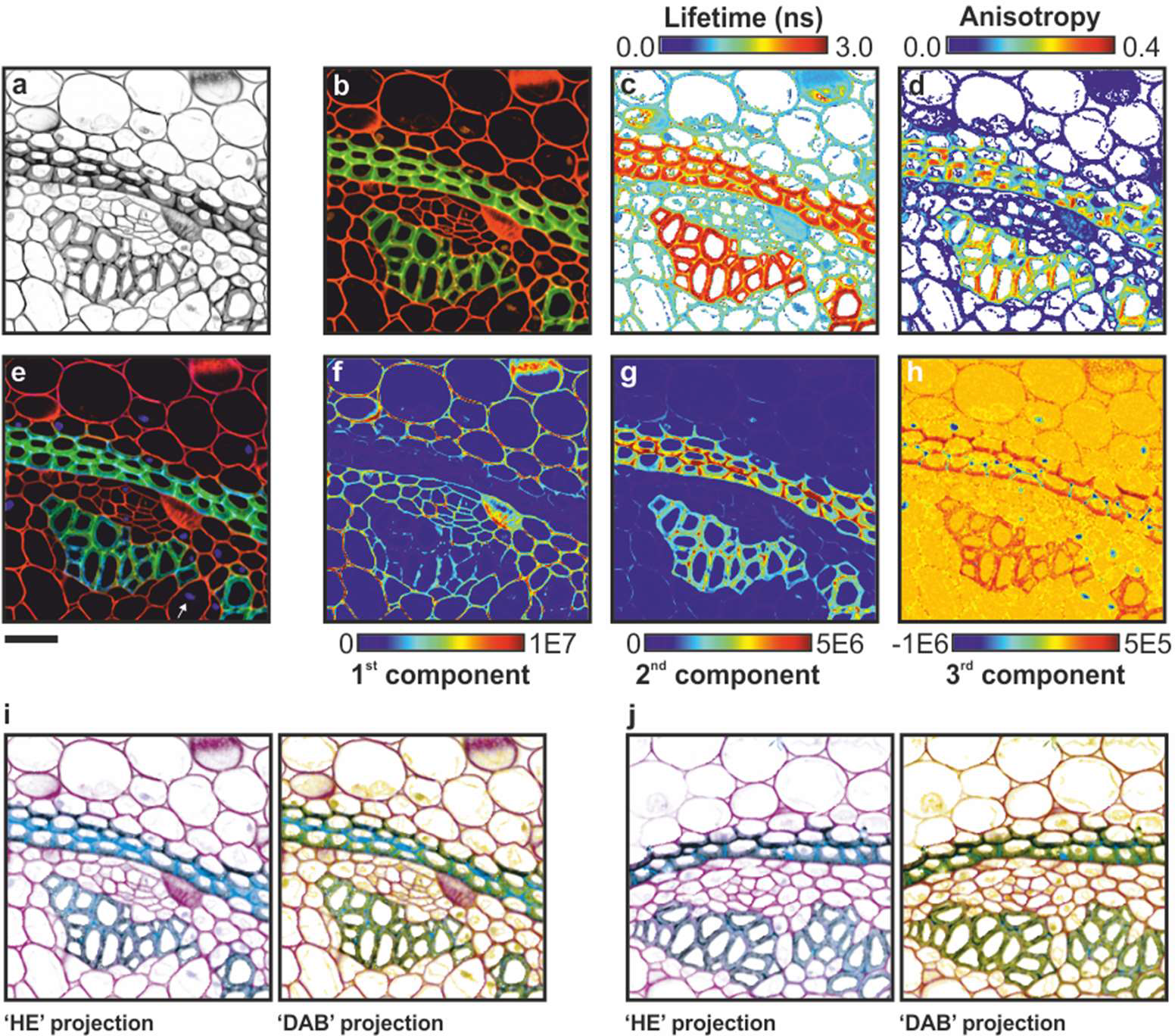
Unmixing by hyper-dimensional imaging microscopy. Images of *Convallaria majalis*, shown as total photon counts (**a**), and projections of HDSS values as true colour representation (**b**), fluorescence lifetime map (**c**), and fluorescence anisotropy map (**d**). Multivariate analysis of HDSS by PCA across the image provide a high contrast RGB composite (**e**) obtained by the overlay of the first three principal components (**f-h**). Scale bar: 40μm; excitation: 800nm.

**Fig. 3i,j** illustrates also how perception matrices can be exploited for possible future applications of HDIM to tissue diagnostics. Principal components can be projected to colour spaces resembling typical counterstains used in histopathology, such as hematoxylin and eosin (HE, **Fig. 3i,j** – left panels, two principal components) and in addition to 3,3′-Diaminobenzidine-like stain (DAB, **Fig. 3i,j** – right panels, three principal components). The full dataset from which **Fig. 3j** was computed is shown in **Supp. Fig. 7**.

Taken together, these observations demonstrate that HDIM provides unprecedented sensing capabilities that can be exploited for contrast enhancement and to better resolve distinct biochemical/photophysical environments.

## Discussion

Previously, we have introduced a generalized concept of resolution in fluorescence microscopy^8^. We have shown that from a theoretical perspective, it is possible to increase information content by increasing the number of independent detection channels, thus enhancing the unmixing capabilities of fluorescence microscopy. Therefore, we hypothesized that the simultaneous detection of orthogonal properties of light, such as fluorescence lifetime, polarization and spectra, could maximize the biochemical resolving power of fluorescence microscopy. To test this hypothesis, we have introduced a new detection paradigm (HDIM), where a time-resolved spectropolarimeter built with off-the-shelf components is utilized to fully resolve the fluorescence emission of specimens. We present here the first experimental evidence that HDIM can, in first instance, provide unprecedented sensing capabilities. Rather than using filters or analysers to generate images during acquisition, HDIM datasets store the full spectroscopic information of a specimen resulting in datasets that can be projected onto spectral, lifetime and polarization dimensions during post-processing. We have also shown that projections can be performed not just on physical features but using statistical tools and perform perception-based projections. We envisage that HDIM will be extremely useful when the optical signatures of a biological phenomenon is unknown, for instance like in tumour imaging where HDIM sensing capabilities can be utilized as an artificial mean of contrast enhancement.

Furthermore, we have shown that – as hypothesized – HDIM results in a significant enhancement of the resolving power in microscopy. Contrary to research aimed to improve the spatial resolution in microscopy which lead to the breakthroughs of modern super-resolution techniques, HDIM exhibits the same spatial resolution of a laser scanning confocal or two-photon microscope; however, HDIM delivers enhanced resolving power to distinguish differences in photophysical properties. Therefore, HDIM can reveal structures that are photophysically distinct but that might be invisible to individual techniques (multi-colour or hyper-spectral imaging, fluorescence lifetime imaging and anisotropy imaging). Therefore, the capability to better resolve distinct emitters should also increase the capability to multiplex a larger number of fluorophores of known characteristics.

With faster and more cost-effective detection technologies being more readily available^9–12^, multiplexed detection technologies could be employed in a range of applications beyond the specialist laboratory, for instance, for multiplexing FRET sensors, for contrast enhancement in label-free tissue imaging or for maximizing the multiplexing capabilities of diagnostic markers in histopathology. We share the software ‘HDIM-toolbox’ aiming to facilitate the community to develop such advanced analytical tools. We propose that heavily multiplexed imaging applications are likely to synergize with emerging technologies such a smart-pixels and deep-learning for machine-vision and a broad range of biomedical applications.

## Materials and Methods

### Microscopy and image analysis

Schematics of the microscope are shown in supporting information (**Supp. Fig. 1 and 2**). Briefly, two photon excitation (TPE) is provided by a tunable femtosecond-pulsed Ti:Sapphire laser (Chameleon Vision 2, Coherent). TPE provides the ideal excitation for nanosecond lived excited state lifetime estimation, high dynamic range for anisotropy measurements and permits the simple separation of the infrared excitation light from ultraviolet-visible fluorescence emission spectra and second harmonic signals. Care should be taken to avoid instabilities of the polarization of the excitation light. The system we developed was built around a Leica SP5 confocal/multi-photon microscope, which uses a periscope formed by a polarization beam splitter (PBS) and a mirror. The PBS works together with a half wave plate to finely tune the excitation power. The poor contrast ratio of a PBS and non-ideal performance of the reflection utilized in the periscope introduced elliptical polarization at the entrance of the microscope which we cleaned-up with a Glan-Thompson polarizer (Newport). HDIM detection was achieved by coupling two grating-based spectrographs (200nm bandwidth) with a polarizer beam splitter. Each spectrograph was equipped with a multi-anode photomultiplier tube and electronics for multi-dimensional time-correlated single photon counting. All TCSPC electronics, detectors and spectrographs were purchased from Becker&Hickl. Equipment for excitation, scanning and detection is commercially available; hardware and software for the integration of these parts have been developed in-house. Th system design is described in **Supporting Figs. 2-3** and can be easily replicated. The HDIM-toolbox is freely available at a GitHub repository (alesposito/HDIM-toolbox) and described in **Supplementary Methods**. At any given image pixel, this setup provides 16 spectral channels over 2 polarization states with arrival times typically histogrammed over 64 time bins. HDIM was calibrated with a laser comb provided by the standard laser lines of the confocal microscope (458nm, 488nm, 514nm, 561nm, 594nm, 633nm) back reflected by the objective lens and with a white light emitting diode which light was scattered by a frosted glass (see **Supplementary Methods**). Typical acquisition times for 256×256 pixels images were within the 1-2 minutes (solutions or *C. majalis* samples). Further details about sample preparation and imaging protocols can be found in **Supplementary Material**. Images were acquired with a HCX PL APO CS 40× 1.25 NA oil objective with a 1.93 zoom thus imaging a field of view of 200μm side.

### Sample preparation

Fluorescent solutions were prepared from ethanol stock solutions of fluorescein (1mM, Fluka Analytical) and rhodamine 6G (5mM, Sigma-Aldrich). Aqueous solutions were prepared with 200μM KCl (Sigma-Aldrich) and a constant ethanol concentration (10% v/v). Fluorophores were quenched by equimolar substitution of KCl for Kl (100μM, Sigma-Aldrich) and their rotational correlation time increased with 65% v/v Glycerol (AnalaR NORMAPUR). The concentrations of the fluorophores were selected to provide a similar brightness with two-photon excitation at 840 nm. *Convallaria majalis* sections stained with Safranin and Fast Green, and mounted were purchased from Leica Microsystems UK (Cat. #As3211).

## Acknowledgements

We acknowledge funding from the Medical Research Council core grants (MC_UU_12022/1 and MC_UU_12022/8), and the EPSRC (EP/F044011/1 and /2). We would like to thank Steve Scotcher, Howard Andrews, Phil Heard, Dave Cattermole and Martin Kyte from the mechanical and electronics workshop at the MRC LMB for their invaluable help with the engineering of our instrumentation. We also would like to thank Bryn Hardwick and Meredith Roberts-Thomson for the initial help with cloning, and Marina Popleteeva for discussion and support. We would like also to thank Leica Microsystems Ltd and Axel Bergmann at Becker&Hickl GmbH for their assistance in integrating electronics and microscopy tools. We acknowledge Prof. Hans Gerritsen (Utrecht University) for long interesting discussions on the topic that led, among other things, to the definition of a clearer names compared to earlier nomenclatures we have used to name early prototypes.

